# Recovery of high-quality assembled genomes via metagenome binning guided with single-cell amplified genomes

**DOI:** 10.1101/2021.01.11.425816

**Authors:** Koji Arikawa, Keigo Ide, Masato Kogawa, Tatusya Saeki, Takuya Yoda, Taruho Endoh, Ayumi Matsuhashi, Haruko Takeyama, Masahito Hosokawa

## Abstract

**Background:** Obtaining high-quality (HQ) reference genomes from microbial communities is crucial for understanding the phylogeny and function of uncultured microbes in complex microbial ecosystems. Despite the improved bioinformatic approaches to generate curated metagenome-assembled genomes (MAGs), existing metagenomic binners often fail to obtain reliable MAGs, and thus, they are nowhere comparable to genomes sequenced from isolates in terms of strain level resolution. Here, we present a single-cell genome-guided metagenome binning (MetaSAG) to reconstruct the strain-resolved genomes from microbial communities at once.

**Results:** MetaSAG employs single-cell amplified genomes (SAGs) generated with microfluidic technology as binning guides to recover improved draft genomes with the metagenomic data. To assess the performance of reconstructing genomes from various microbial communities, we compared MetaSAG with four conventional metagenomic binners using a cell mock community, human gut microbiota, and skin microbiota samples. MetaSAG showed precise contig binning and higher recovery rates (>97%) of rRNA and plasmids compared to conventional binners in genome reconstruction from the cell mock community. In human microbiota samples, MetaSAG recovered the largest number of genomes with a total of 103 gut microbial genomes (21 HQ and 65 showed >90% completeness) and 45 skin microbial genomes (10 HQ and 40 showed >90% completeness), respectively. Conventional binners recovered one *Staphylococcus hominis* genome, whereas MetaSAG recovered two *S. hominis* genomes from the identical skin microbiota sample. Single-cell sequencing indicated that these *S. hominis* genomes clearly derived from two distinct strains harboring specifically different plasmids. We found that all conventional *S. hominis* MAGs had substantial lack or excess of the genome sequences and contamination of other *Staphylococcus* bacteria (*S. epidermidis)*.

**Conclusions:** MetaSAG enabled us to obtain the strain-resolved genomes in the mock community and human microbiota samples by assigning metagenomic sequences correctly and covering both highly conserved genes such as rRNA genes and unique extrachromosomal elements, including plasmids. MetaSAG will provide HQ genomes that are difficult to obtain with metagenomic analyses alone and will facilitate the understanding of microbial ecosystems by elucidating detailed metabolic pathways and horizontal gene transfer networks through microbial genomes. MetaSAG is available at https://github.com/kojiari/metasag.

## Background

The accumulation of reference genomes from microbes has provided insight into the ecology and evolution of environmental and host-associated microbiomes. The golden standard for microbial genome sequencing has been to culture specific strains and sequence extracted DNA[1–3]. Recently, metagenomic analysis, which combines direct extraction of genomic DNA from the microbial community with *in silico* reconstruction of each microbial genome sequence from massive sequenced reads, has attracted much attention. A growing number of metagenome-assembled genomes (MAGs) are asserting our understanding of microbial diversities in various environments[4–9].

In a metagenomic approach, genome reconstruction is performed in two steps: (1) assembly of fragmented genome sequences to contigs and (2) binning contigs into lineages as bins. Current state-of-the-art binners rely on nucleotide compositional information such as tetranucleotide frequency, GC content, or sequence coverage[10–12]. However, these tools demonstrate different performances and produce different MAGs including incomplete bins and multi-species composite bins[13]. Composite genomes that aggregate sequences originating from multiple distinct species or strains can yield misleading insights if they are registered as single genomes in the reference database[14]. To solve these problems, several approaches combine and curate the result of multiple binners to generate a large number of high-quality (HQ) genomes[13,15,16]. However, in the real samples, it is difficult to verify the certainty of the binning results because there are numerous microbes without the reference genome and the proportion of microbial species richness among them is unknown.

Single-cell genomics is an alternative approach that enables culture-independent sequencing of microbial genomes[17]. In contrast to metagenomics, single-cell genomics does not require microbial population clonality but instead recovers genome sequences from individual cells. In single-cell genomics, the DNA amplification process often causes amplification biases and incompleteness in genome sequences. Therefore, co-assembly of individual single-cell sequencing data is generally required to compensate for the gaps and errors in each SAG sequence[18]. However, most SAGs generally have low completeness, and even with co-assembly, produced shortly fragmented contigs, rarely covering their entire genomic area.

Metagenomics assesses the genomes of all microbes present in a sample, whereas single-cell genomics reveals individual genomes. Therefore, it has been suggested that integrating the two can compensate for each of their specific shortcomings[19–21]. However, no efforts have been made to acquire multiple draft genomes of the human microbiota using this hybrid approach. Moreover, its advantages over the conventional metagenomics binning have not been verified. In this study, we developed a novel metagenome binning guided with single-cell amplified genomes (MetaSAG) to recover at once HQ genomes of multiple bacterial strains from the microbial community. We used microfluidic technology-aided approaches to obtain a large number of single-cell amplified genomes (SAGs) for guided binning[22,23]. Mock community and human microbiota samples were tested to compare the binning accuracy and the number of HQ genomes between conventional binners and MetaSAG. We also investigated the integration of single-cell genomes with metagenomes to acquire strain-resolved genomes and to validate the presence of aggregate sequences originating from multiple distinct species in metagenomic bins

## Results

### Overview of the single-cell genome-guided metagenome binning (MetaSAG)

Conventional metagenomic phylogenetic classification tools[24,25] and conventional metagenomic binners[10–12] have difficulty in allocating contigs to bins from complex microbial communities in the absence of known microbial genome information as teaching data for classifying closely related species or strains. Our MetaSAG tool uses single-cell amplified genomes, which are newly produced in the same sample, as teaching data for metagenome binning (Fig. 1). The SAGs of uncultured microbes serve as ideal references for metagenome binning from the reference-lack microbial community. These SAGs were obtained using the SAG-gel platform[22,26], which enables obtaining the contamination-less SAGs in a high throughput manner with the aid of microfluidic droplet format. Multispecies SAGs obtained by assembly from each single-cell genome are grouped into individual strains using the ccSAG method[18]. Composite SAGs (CoSAGs) are constructed by re-assembling (co-assembling) single-cell reads (SRs) recognized as identical strains. Based on genome completeness (>50%) and contamination (10%), non-redundant SAGs (nrSAGs) are collected for use as binning references. Besides, metagenomic reads (MRs) are obtained from the same sample and are assembled into metagenomic assembled contigs (MAs). The contigs in nrSAGs are mapped to the contigs in MA to allocate contigs in MAs to single cell-guided bins (sgBins). Finally, the paired nrSAGs and sgBins at the strain level are merged to fill in the gaps for each other and extend the contig length as single-cell-guided MAGs (sgMAGs) or metagenome-guided SAGs (mgSAGs).

**Fig. 1.**
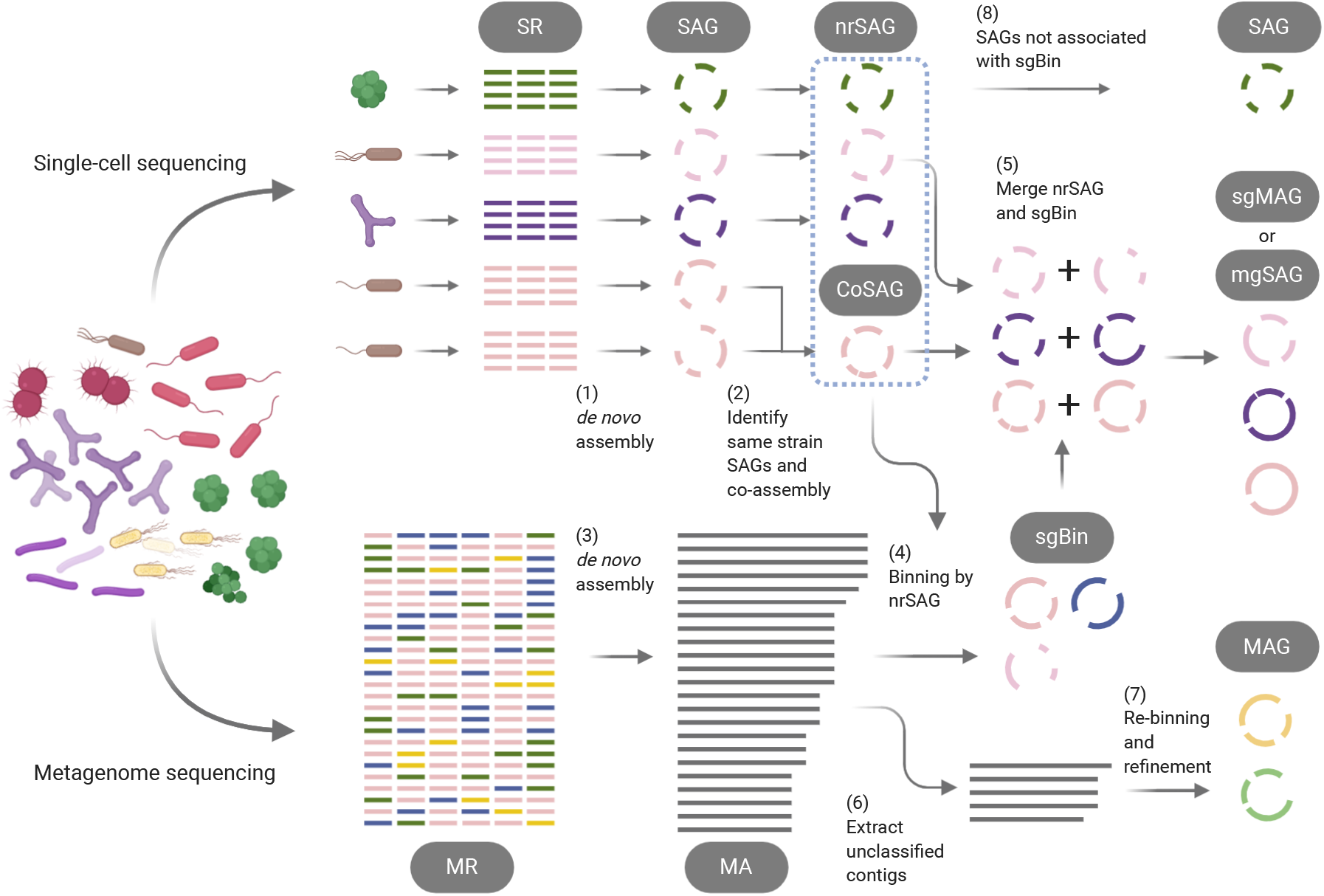
Overview of metagenome binning guided with single-cell amplified genomes (MetaSAG) workflow. Single-cell sequencing reads (SRs) and metagenomic sequencing reads (MRs) are obtained from the same microbial community. (1) *De novo* assembly of each SR to a single-cell amplified genome (SAG). (2) SAGs of the same strain are identified into the group and co-assembled into composite SAG (CoSAG). (3) *De novo* assembly of MRs to metagenome-assembled contigs (MAs). (4) MA is classified to single-cell genome-guided bin (sgBin) by mapping MA on non-redundant SAG (nrSAG). (5) Paired nrSAGs and sgBins are merged to single-cell genome-guided MAG (sgMAG) or metagenome-guided SAG (mgSAG). (6) Unbinned contigs in MA are extracted and subsequently (7) re-binned and refined by conventional metagenomic binning and refinement tools. Four types of draft genomes (SAG, sgMAG, mgSAG, and MAG) are finally acquired.

### Evaluation of single-cell genome and metagenome assemblies

To confirm sequence accuracy in nrSAGs and MAs, single-cell genomic and metagenomic sequencings were performed with the same cell mock community containing 15 different bacteria (Additional file 1: Table S1). In total, we obtained 48 SRs and one MR with total read lengths of 3.9 Gb and 2.6 Gb, respectively (Additional file 2: Table S2).

Following the assembly, 15 nrSAGs were obtained, which covered all species in the mock community. From SAG to CoSAG according to taxonomy identification (Additional file 3: Table S3), the average completion rates improved from 33.5% to 66.6%, with low contamination rates of 0.3% and 0.76%, respectively (Additional file 4: Table S4). In 14 nrSAGs, approximately ≥98.5% of the total length of each was correctly mapped to reference genomes. In Mock-C00006 (*Lactobacillus delbrueckii*), some contigs (8.5% of the total length) were mapped to other microbial genomes. The original SAGs were obtained from physically isolated single-cells in gel capsules[22]; however free DNA was randomly captured and subsequently amplified simultaneously. The unmapped contigs could have been derived from these free DNA fragments. Alternatively, we confirmed that 1008 contigs of total 1016 MA contigs were mapped to single reference genomes (Additional file 5: Fig. S1). In addition, there were no 16S rRNA gene sequences for *Bacteroides uniformis* and *Escherichia coli* in MA, while all nrSAGs remained individual 16S rRNA sequences (Additional file 5: Fig. S2). Overall, both the nrSAGs and the MA showed a high sequence accuracy as high identity corresponding to reference genomes. Thus, we considered that the metagenomic binning step was crucial for reconstructing each genome from the MA.

### Comparison of the characteristics of single-cell-guided bins with conventional bins

We investigated the characteristics of bins collected by MetaSAG and conventional binners (Fig. 2). Three binners, CONCOCT[10], MaxBin2[12], and MetaBAT2[11], were used to construct bins, and DAS_Tool[13] was subsequently used to obtain refined bins. Based on 15 reference genomes (Additional file 1: Table S1), we assessed the taxa of each bin and estimated the total sizes of the contigs incorrectly binned to different bacterial bins and contigs unbinned to any reference genomes (Fig. 2a). The binner with the smallest incorrect binned contig was MetaSAG with 20 kbp, followed by MetaBAT2 with 181 kbp. The binner with the smallest unbinned contig was CONCOCT (1 kbp), while the unbinned contig length was 892 kbp in MetaSAG. Total lengths of contigs unbinned into target sgBins were inversely correlated with the guide SAG completeness (Fig. 2b), suggesting that the SAG completeness strengthens the adequacy of the contig allocation to the bin.

**Fig. 2.**
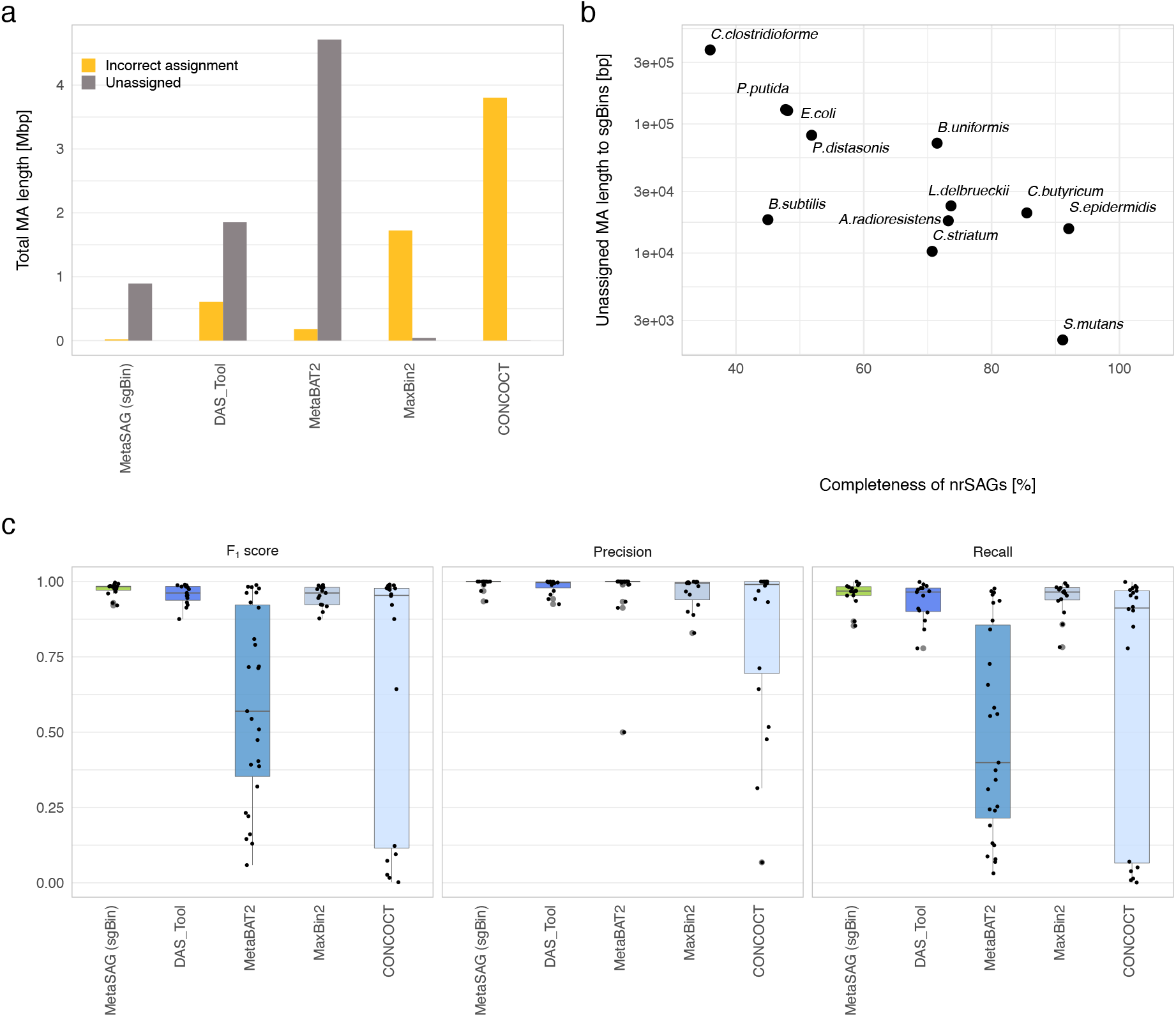
Precision of single-cell-guided binning of metagenome assembly from a mock microbial community of 15 bacteria. (a) Total length of contigs incorrectly binned to different species and unbinned metagenome assembled contigs (MAs). (b) Correlation between completeness of non-redundant single-cell amplified genomes (nrSAGs) and total length of contigs unbinned to the target single-cell genome-guided bin (sgBin). (c) The plots of F1 scores, precision, and recall of all reported bins (center line, median; box limits, upper and lower quartiles; whiskers, minimum or maximum values between upper and lower quartiles that are extended 1.5 times the interquartile region). Individual values are represented as dots.

We also calculated F1 scores, a harmonic mean of precision and recall, to evaluate the accuracy of bins construction against true reference genomes (Fig. 2c). The precision depends on the less false-positive contig that is incorrectly allocated in the bin. Although the ability to force contigs into bins helps improve completeness, it carries the risk of inclusion of artificial sequences as false-positive contigs and increases contamination rates. MetaSAG demonstrated high-precision bins against all 15 corresponding references. The high-precision bin (F1 score >0.9) for MetaSAG, DAS_Tool, MetaBAT2, MaxBin2, and CONCOCT were 15, 14, 8, 13, and 12, respectively, while all binners except MetaBAT2 had equally high precision values. Alternatively, the recall value depends on the true completeness of the bacterial genome. MetaSAG demonstrated the highest F1 scores among all reference genomes owing to the highest recall value. In this test, SAG qualities were limited to low-quality (LQ) to medium-quality (MQ), which were not the best conditions to guide binning; however, it was still remarkably clear that MetaSAG had the best binning accuracy. Thus, the single-cell guided binning approach of MetaSAG helps accurate and efficient allocation of contigs into different bacterial genomes compared to conventional binners.

### Effectiveness to merge nrSAGs and sgBins

The merging of paired nrSAGs and sgBins into sgMAG or mgSAG improved several genome assembly quality metrics such as completeness and N50 in multiple microbial communities including human gut microbiota, and human skin microbiota (Fig. 3ab). Although the completeness of either nrSAG or sgBin was low (average: 74.5%), that of sgMAG and mgSAG was much improved (average: 93.6%) (Fig. 3a). N50 metrics of most nrSAGs (average: 48.2 kb) improved after merging nrSAG and sgBin (average: 87.7 kb), except in the case of low completeness of sgBins (Fig. 3b). Low completeness of sgBins occurred often, particularly in skin microbiota (average completeness: 23.1%). This may be due to the inability of metagenomic data to produce qualified MAs owing to some interfering factors, such as human DNA contamination. Thus, to recover sgBins with high completeness, it is necessary to increase the MA mapping rate by improving its assembly accuracy and by increasing SAGs repertoire corresponding to MAs. In addition, rRNA and tRNA gene sequences were often compensated from nrSAGs (Recovery rate of rRNA: 5S: >53.1%, 16S: >94.1%, 23S: >98.5% in nrSAGs; and 5S: >7.5%, 16S: >13.4%, 23S: >14.9% in sgBins)(Fig. 3cb), thus merging of nrSAGs and sgBin is extremely important for incorporating phylogenetic information of draft genomes.

**Fig. 3.**
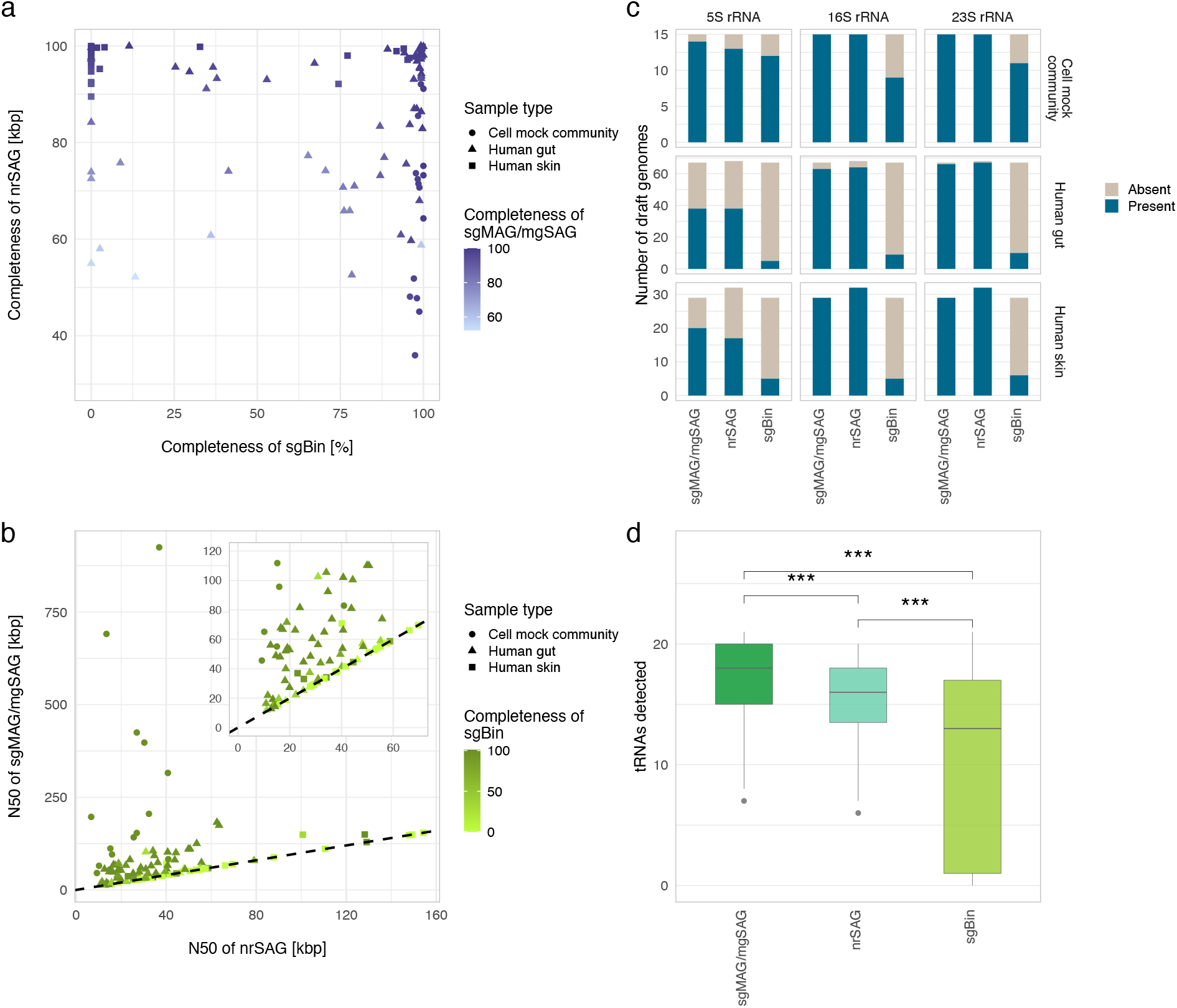
Assembly quality metrics of single-cell-guided metagenome-assembled genome (sgMAG) and metagenome-guided single-cell amplified genome (mgSAG). All data were collected from a mock community containing 15 bacteria, three human fecal samples, and three human skin swab samples. (a) Scatter plot of completeness of non-redundant single-cell amplified genome (nrSAGs) versus single-cell genome guided bins (sgBins) corresponding to all medium-quality (MQ) and high-quality (HQ) sgMAGs and mgSAGs. (b) Relationship between N50s of nrSAG and of sgMAG or mgSAG. Numbers of rRNA genes (c) and tRNA genes (d) in draft genomes produced in MetaSAG workflow (center line, median; box limits, upper and lower quartiles; whiskers, minimum or maximum values between upper and lower quartiles that are extended 1.5 times the interquartile region, Wilcoxon rank sum test ***: *p* <0.001).

### Recovery of HQ draft genomes from multiple microbial communities using MetaSAG

We assessed the quality of all draft genomes according to the Genomic Standards Consortium[9]. From the mock community sample, MetaSAG, DAS_Tool, and MaxBin2 constructed MAGs corresponding to 15 reference genomes, whereas MetaBAT2 and CONCOCT constructed more than the expected 15 MAGs, including several LQ MAGs (Fig. 4a). Thus, the risk of creating unreliable MAGs must also be deliberated when considering conventional binners. MetaSAG uses nrSAG taxonomy to identify representative species and to extract contigs in MA necessary for binning, such that the risk of producing artificial MAGs that cannot be present in actual samples is diminished. Regarding draft genome quality, MetaSAG produced a total of 13 HQ draft genomes, with better accuracy than other binners (Fig. 4a). For non-chromosomal elements, all plasmid sequences existed in MA; however, these were lost in the plasmid-containing bacterial genomes after binning (Additional file 5: Fig. S3). MetaSAG demonstrated constant and higher plasmid coverage (97.2%) than other binners (50.6-74.5%) in five bacteria.

**Fig. 4.**
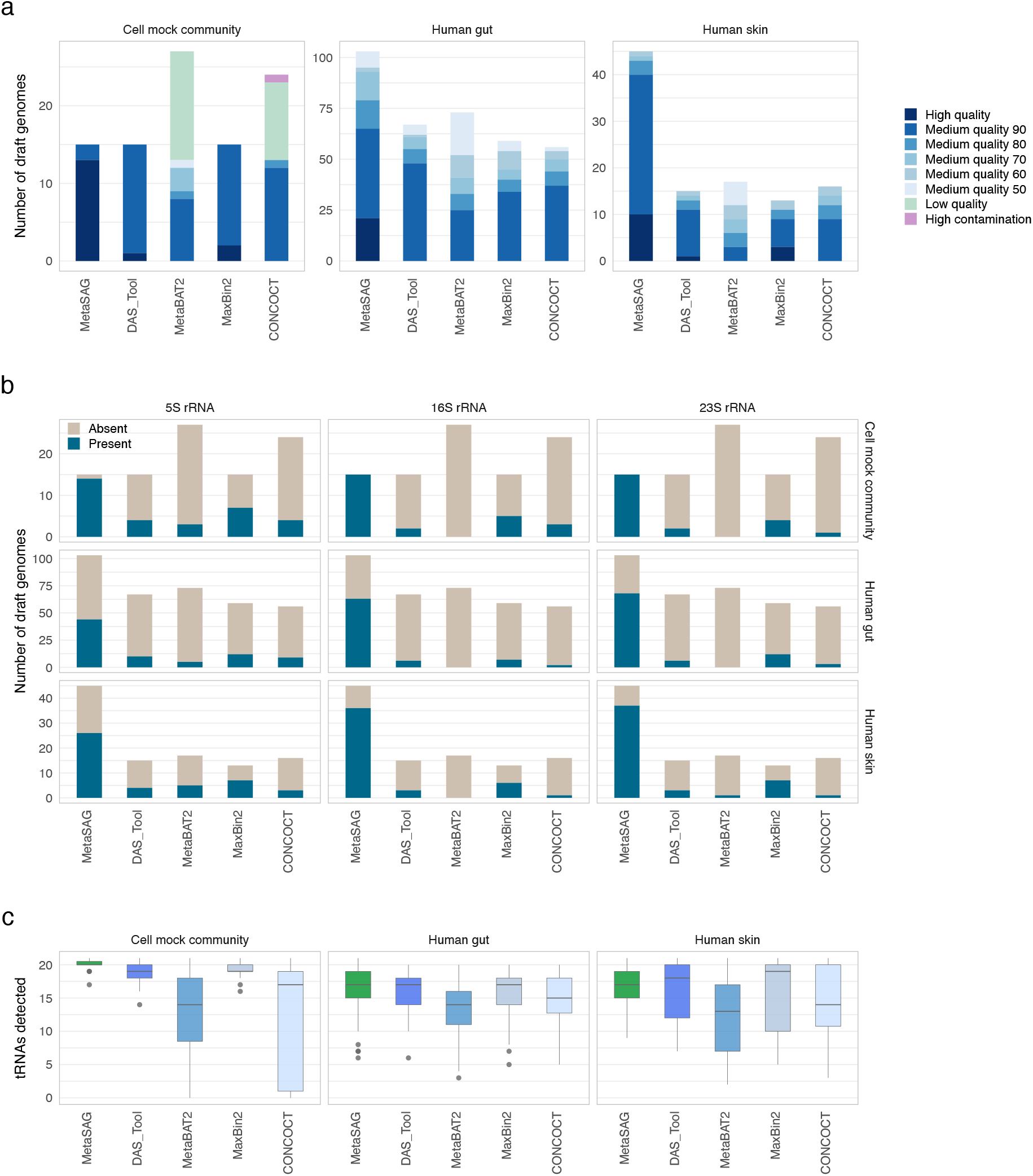
Draft genomes reconstructed from the cell mock community and human microbiota samples. All data were collected from a mock community containing 15 bacteria, three human fecal samples, and three human skin swab samples. (a) The number of reconstructed genomes per method. Human gut and skin data show medium-quality (MQ) and high-quality (HQ) genomes only. Number of rRNA genes (b) and tRNA genes (c) in draft genomes produced with MetaSAG and other tools (center line, median; box limits, upper and lower quartiles; whiskers, minimum or maximum values between upper and lower quartiles that are extended 1.5 times the interquartile region).

To evaluate the performance of MetaSAG in human gut and skin microbiota, three SR sets (each 96 SR, 100 Mb/SR) and three MRs (each 6 Gb) were obtained, and the assemblies were binned with MetaSAG and other binners. Here, MQ and HQ draft genomes were considered for comparison. MetaSAG was able to construct the largest number of genomes with a total of 103 (21 HQ) and 45 (10 HQ) genomes from the gut and skin, respectively (Fig. 4a and Additional file 6: Table S5). In gut microbiota, none of the HQ genomes were constructed in other binners. Although there were draft genomes that satisfied a completeness >90% and contamination <5% in conventional binners, difficulty in recovery of rRNA and tRNA sequences were clearly indicated (Fig.4 bc). MetaSAG demonstrated consistent high performance in the recovery of rRNA (5S: >42.7%, 16S: >61.2%, and 23S: >66.0%) and tRNA (average: 17.3 ± 2.9) in each microbial sample. MetaSAG used a large number of sequencing reads by incorporating single-cell genomics and metagenomics; however, trends were unchanged, even when the read number used for other binners was equal to that when MetaSAG was used (Additional file 5: Fig. S4).

### Coverage of MetaSAG-produced draft genomes against bacterial diversity

To determine the extent to which the constructed genome covered all metagenomic sequence fractions, MRs were mapped to their respective genomes and mapping rates were calculated. In MAGs constructed by MaxBin2 and CONCOCT, >90% of MRs were mapped (Fig. 5a). These high mapping rates were considered owing to their algorithm trends of unbinned contig reduction (Fig. 2a). The MR mapping rates in MetaSAG were in the middle of all binners, ranging from 78.9% to 89.5% for gut microbiota and 91.3% to 95.6% for skin microbiota. Regarding the bacterial diversity, MetaSAG detected more bacterial genomes than other binners, with 54 genera in gut microbiota (Fig. 5b) and nine genera in skin microbiota (Fig. 5c). We considered that the metagenomic coverage of MetaSAG could be improved by increasing the number of obtained SAGs from the same samples and the number of detected taxa.

**Fig. 5.**
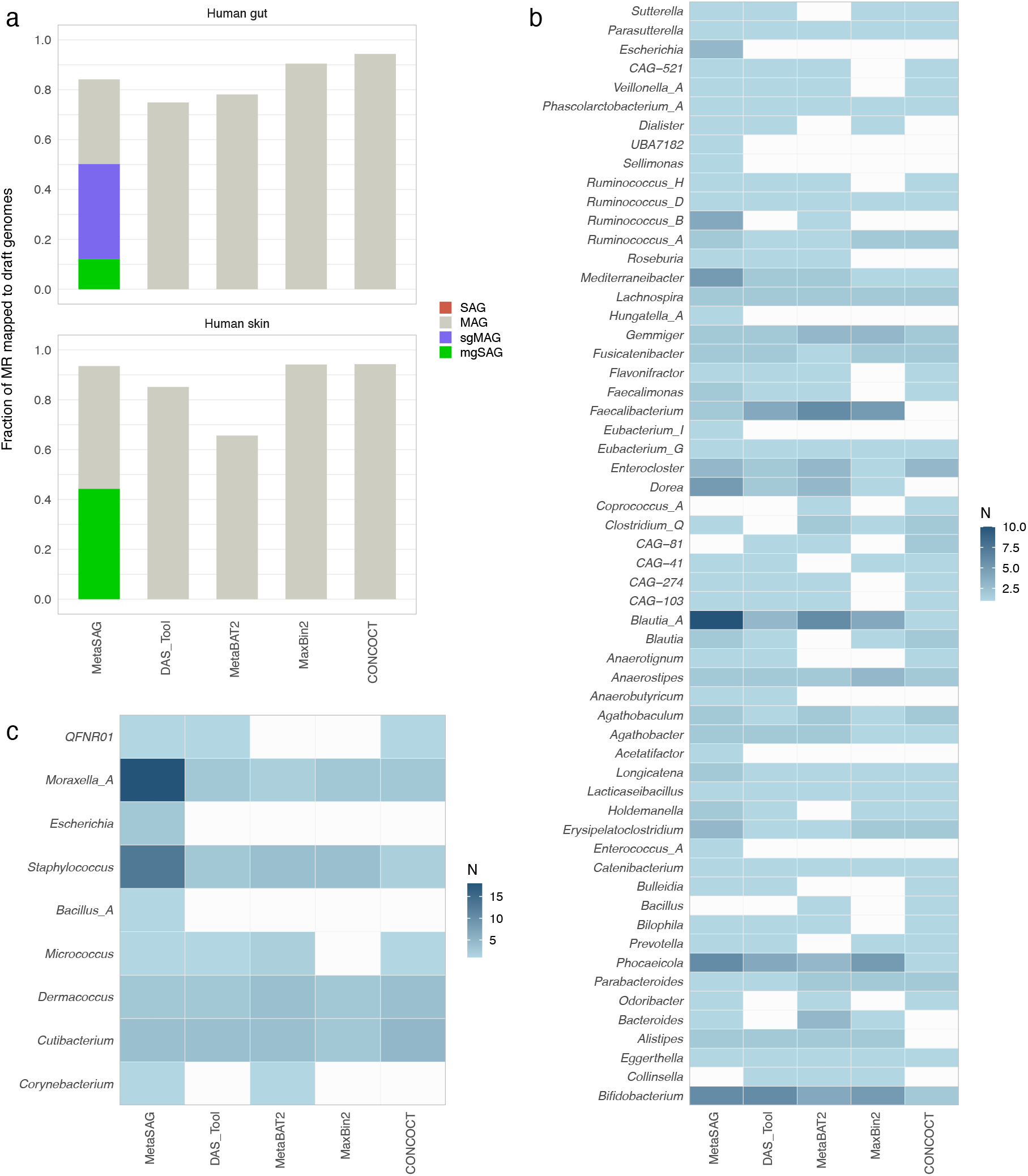
Diversity of draft genomes reconstructed by MetaSAG. (a) Fraction of metagenomic reads against draft genomes constructed by each binner. MetaSAG shows a fraction of metagenomic reads against four types of draft genomes. The number of draft genomes acquired from (b) human gut and (c) human skin are collapsed by genus assigned with GTDB-Tk.

### Strain-resolved genome analysis based on single-cell genomes toward revealing intra-species diversities

Accurate genomic classification of closely related species and subspecies from the microbial community is important and is required to discuss intra-species diversity. Therefore, to assess separation accuracy of closely related genomes, we assessed the correspondence between MAG and SAG sequences of the same species.

In skin microbiota, conventional metagenome binner yielded one *Staphylococcus hominis* MAG, whereas MetaSAG yielded two *S. hominis* strain mgSAGs (*S. hominis* BBMGS-S01-101 and *S. hominis* BBMGS-S01-100). We considered that conventional MAG had difficulty in binning contigs to two different strains in the sample. To confirm further details, we calculated the average nucleotide identity (ANI) of the two strain genomes obtained by MetaSAG and other binners against the original SAGs (Fig. 6a) and we confirmed all ANI showed >97% identities. We found that while the presence of two strains is evident at the single-cell level, only MetaSAG was able to output strain-resolved genomes, and the binner produced MAGs which demonstrated increased similarity to only one strain. Notably, we found plasmids in conventional MAGs; however, plasmid assignment to mgSAGs clearly indicated that these two strains had specifically different plasmids (Fig. 6b). Thus, MetaSAG will aid strain-resolved binning and plasmid-host allocation, resulting in an understanding of intra-species diversities and linking mobile gene elements to hosts.

**Fig. 6.**
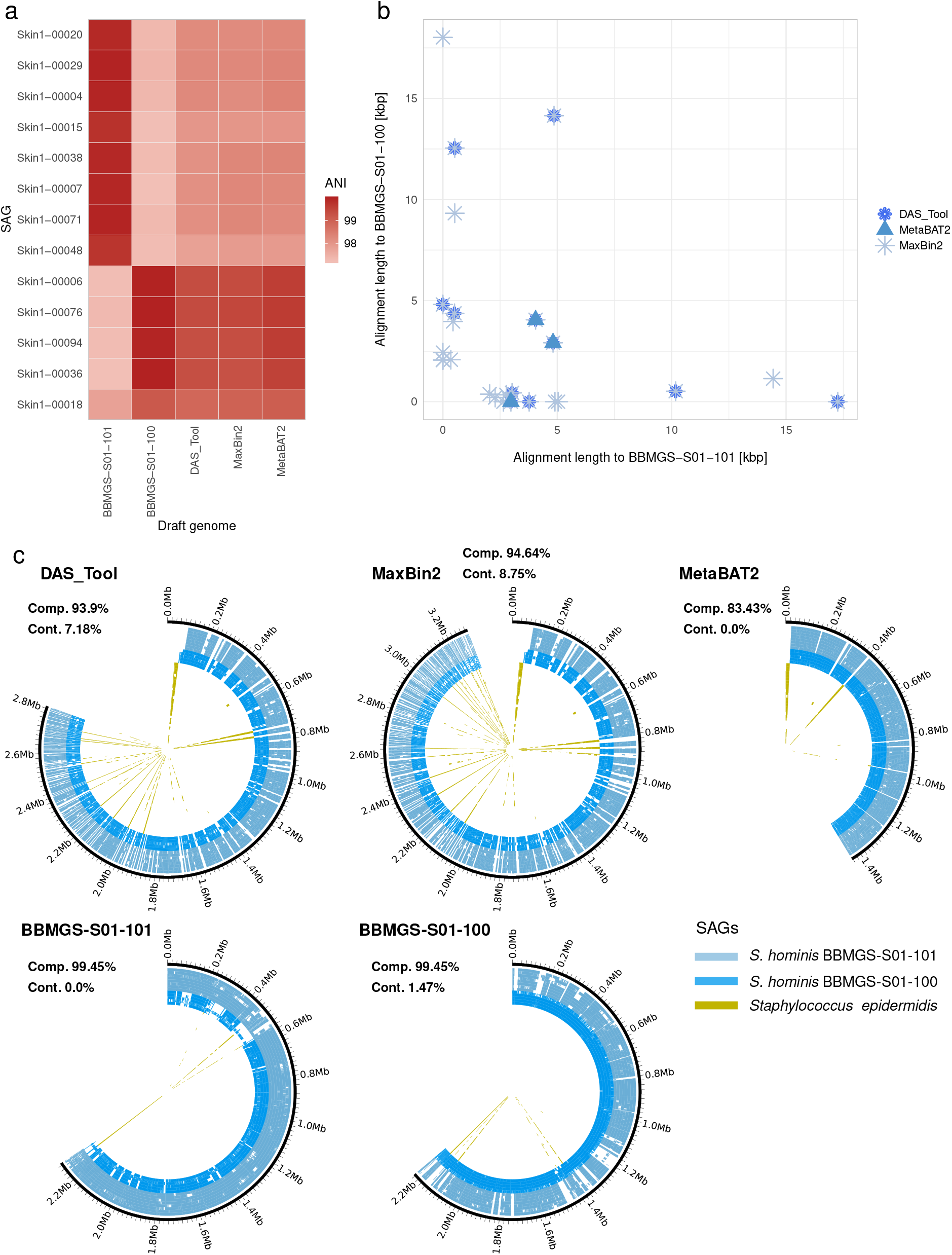
Strain-resolved analysis of skin microbes for host-plasmid linking and detection of interspecies chimeric sequences. (a) Mean pairwise genomic similarities between *Staphylococcus hominis* draft genomes obtained with MetaSAG (*S. hominis* BBMGS-S01-101 and *S. hominis* BBMGS-S01-100), and other binners. (b) The scatter plot shows the length of plasmid contigs assigned to BBMGS-S01-101 and BBMGS-S01-100. Different plot symbols show contigs obtained with different binners. (c) Interspecies chimeric sequence detection by alignment of *S. hominis* MAGs derived with conventional binners (yellow), and *S. hominis* BBMGS-S01-101 (light blue) and *S. hominis* BBMGS-S01-100 (blue). The outermost circles show draft genomes and the inner circles show the result of mapping individual SAGs, which belong to the same genus of *Staphylococcus*, to the draft genome.

### Validation of aggregate sequences originating from multiple distinct species

SAG can be used as a self-check reference to evaluate the appropriateness of conventional MAG binning results, and it may also be possible to remove unsuitable contigs such as aggregate sequences from multiple species. A simple way to detect incorrect sequences in MAG is to map corresponding SAG sequences to MAGs (Fig. 6c). For *S. hominis* obtained from human skin microbiota, we screened SAG sequences which were mapped on MAGs obtained with conventional binners. This result clearly indicated that *S. hominis* MAGs showed different genome sizes between different binners (1.4 to 3.2 MB) while they showed high completeness (83% to 94%), suggesting substantial lack or excess of the genome sequence, and some sequences from the closed *Staphylococcus* genus (*S. epidermidis*) were contaminated in all MAGs (55.6-146.8kb). In particular, we found that the longest contaminated contig (44 Kb) in MAGs of DAS_tool and MaxBin2 showed homology (identity 98.5%) to pSE2 plasmid of *S. epidermidis* (CP066374). The genome sizes of publicly available *S. hominis* isolate genomes are 2.1 to 2.3 Mb and are similar to the draft genome obtained with MetaSAG. BBMGS-S01-101 and BBMGS-S01-100 exhibited some common sequences between *S. hominis* and *S. epidermidis*; however, there were no obvious interspecies aggregate sequences. Using SAGs as references, contigs that have been erroneously removed or included by conventional binners can be correctly assigned, suggesting that even uncultured bacterial genomes can be validated for showing their reliability at the strain-level.

## Discussion

HQ reference genomes are essential for understanding the phylogeny and function of uncultured microbes in complex microbial ecosystems. In a changing environment, microbes acquire adaptive evolution through repeated genetic mutations and horizontal transfer, etc.[27–30]. To best understand the connections between microbial communities and their habitats is to recover genomes from the communities themselves, rather than referring to genomes of closely related bacteria isolated from different environments.

Despite the cell mock community being a simple sample consisting of 15 different bacteria, the occurrence of false-positive contigs in the conventional MAG suggested the requirement for careful selection of the metagenomic binner depends on the presence of conserved genes and consistency of nucleotide composition. As reported previously[13,15], the tool that utilizes the bin refinement strategy demonstrated high accuracy, which was in agreement with MAG and reference genomes. These tools utilize multiple binners to generate various combinations of bins for reference to each other from single or multiple metagenomics data. Alternatively, MetaSAG generates self-references from the same sample at the single-cell level and they are subsequently guided to bin metagenomic contigs for genome reconstruction. MetaSAG enabled us to obtain the highest qualities in draft genomes, both in the mock and human microbiota samples, by assigning metagenomic sequences in correct bins, as well as by filling the gap in highly common sequences, such as rRNA genes, and linking the host with extrachromosomal elements, such as plasmids. The integration of metagenomics and single-cell genomics has been used to improve genome recovery from environmental bacteria. It was previously reported that metagenomic reads can be used to fill in gaps in SAGs[19], or that SAGs can be used as scaffolds for MAGs[20]. However, the number of constructed genomes was limited, and no tool has been developed to obtain multispecies genomes at once, which is mostly due to the lack of technology that provides good quality SAGs as binning guides. In MetaSAG, the qualities of SAGs obtained by our SAG-gel technology[22] were sufficiently high to prevent false-positive contigs in supervised contig identification. In addition, merging of SAGs with the metagenomic bin aided recovery of rRNA and tRNA sequences, which were frequently lacking in the MAGs obtained by conventional binners. This advantage overcomes the incompleteness of phylogenetic information contained in conventional metagenomic bins, suggesting that this technology can be used to move forward from conventional microbial profiling using 16S rRNA gene amplicon sequencing to metabolic function analysis referring to novel genomes.

One of the challenges of MetaSAG is the difficulty in obtaining genome sequences beyond the number of SAGs acquired in advance. To obtain genomes from samples of high microbial diversity or to obtain genomes of rare microbes, it is necessary to obtain a large number of SAGs or to selectively obtain SAGs of the desired taxa. In this study, we recruited the SAGs with the completeness >20% to produce CoSAG with the completeness >50%. To improve the genome number, the approaches are considered to accumulate massive LQ SAGs with low sequencing efforts to produce nrSAG which covers a broad microbial spectrum, or target single-cell genome sequencing with species enrichment techniques[17,31,32]. Another issue with MetaSAG is that it only allows allocation to a single sgBin per contig for binning using nrSAG as a guide. Under this binning condition, if there are multiple bacterial strains with extremely similar sequences, the assignment of MA contig to sgBin may not be fulfilled in any of the strain genomes. Nonetheless, the implementation of contig assignment to multiple sgBin requires careful consideration owing to the complexity of the computational process and the possibility of producing interspecies aggregate sequences. We recommend using mgSAG, where the completeness of the SAG itself is increased and used as primary data, and the metagenome is used as supplementary information. This procedure allows us to obtain strain-resolved genomes and observe differences among strains, taking advantage of the resolution of SAGs.

MetaSAG has the ability to control the SAG integration level by adjusting parameters. It is possible to construct representative sequences for each taxonomy rank by appropriately setting single copy marker gene homology, ANI, and tetranucleotide frequency, which are parameters used for SAG integration to CoSAG. These SAGs can be utilized as reference genome sequences against which resulting MAGs are checked for harboring interspecies aggregate sequences. Verification of the reliability of MAGs is critical because composite genomes that aggregate sequences from several different populations can provide misleading insights when treated and reported as a single genome. By using MetaSAG, if biological samples that are the source of metagenomic data are properly stored and new single-cell data can be obtained, we will be able to increase the accuracy of acquired data curation and MAG by obtaining new single-cell genomes. Besides, MetaSAG can subdivide genomes of individual strains, even for species that cannot be divided into strain levels by metagenomic bins. Single-cell based strain-resolved genome analysis will contribute to our understanding of intraspecies diversity and distribution of non-chromosomal elements[29,33–35].

## Conclusion

In conclusion, MetaSAG can integrate SAG and MAG to reconstruct qualified microbial genomes and control their binning resolution based on the numbers and classification of SAGs. Since it can provide reliable HQ genomes from a variety of microbial communities, it will represent a powerful tool to support microbial research that requires reference genome expansion and strain-resolved analysis toward understanding microbial association to the host or environment. Thus, MetaSAG is highly scalable and can be applied to reuse previously acquired metagenomics data and single-cell genomics tools to be developed.

## Methods

### Experimental design and sample collection

Fresh feces were collected by subjects in 15 mL vials containing 3 mL GuSCN solution (TechnoSuruga Laboratory Co., Ltd,) and stored for 2 d maximum, prior to DNA extraction and single-cell encapsulation in droplets.

Skin bacterial samples were collected and placed in Dulbecco’s phosphate-buffered saline (DPBS) by swabbing the surface of facial skin using sterile cotton applicators (Nissui Pharmaceutical Co., Ltd) pre-moistened with DPBS by subjects and were stored at room temperature for 2 d maximum, prior to DNA extraction and single-cell genome amplification.

The mock microbial community (Cell-Mock-001) was obtained from the National Institute of Technology and Evaluation Biological Resource Center, Japan. This mock microbial community was composed of 15 bacterial strains detected in various environments (intestinal, oral, skin, and natural environment) as follows: *Bacteroides uniformis, Bifidobacterium pseudocatenulatum, Clostridium clostridioforme, Cutibacterium acnes subsp. acnes, Escherichia coli* K-12, *Parabacteroides distasonis, Staphylococcus epidermidis, Streptococcus mutans, Acinetobacter radioresistance, Comamonas terrigenous, Bacillus subtilis subsp. subtilis, Clostridium butyricum, Corynebacterium striatum, Lactobacillus delbrueckii subsp. delbrueckii*, and *Pseudomonas putida*.

### Single-cell genome sequencing with SAG-gel

Single-cell genome sequencing was performed with single-cell whole genome amplification (WGA) using the SAG-gel platform according to our previous reports[22,26]. Following homogenization of human feces in GuSCN solution (500 μL), the supernatant was recovered by centrifugation at 2000 ×*g* for 30 s, followed by filtration through 35-μm nylon mesh and centrifugation at 8,000 ×*g* for 5 min. The resulting cell pellets were suspended in PBS, washed twice at 8,000 ×*g* for 5 min. Skin swab samples in DPBS were processed in the same manner except for homogenization.

Prior to single-cell encapsulation, cell suspensions were adjusted to 0.1 cells/droplets in 1.5% agarose in PBS to prevent encapsulation of multiple cells in single droplets. Using the droplet generator (On-chip Biotechnologies Co., Ltd.), single microbial cells were encapsulated in droplets and collected in a 1.5-mL tube, which was chilled on ice for 15 min to form the gel matrix. Following solidification, collected droplets were broken with 1H,1H,2H,2H-perfluoro-1-octanol (Sigma-Aldrich) to collect beads. Thereafter, the gel beads were washed with 500 μL acetone (Sigma-Aldrich), and the solution was mixed vigorously and centrifuged. The acetone supernatant was removed, 500 μL isopropanol (Sigma-Aldrich) was added, and the solution was mixed vigorously and centrifuged. The isopropanol supernatant was removed, and the gel beads were washed three times with 500 μL DPBS.

Thereafter, individual cells in beads were lysed by submerging the gel beads in lysis solutions: first, 50 U/μL Ready-Lyse Lysozyme Solution (Epicentre), 2 U/mL Zymolyase (Zymo research), 22 U/mL lysostaphin (MERCK), and 250 U/mL mutanolysin (MERCK) in DPBS at 37 °C overnight; second, 0.5 mg/mL achromopeptidase (FUJIFILM Wako Chemicals) in PBS at 37 °C for 8 h; and third, 1 mg/mL Proteinase K (Promega) with 0.5% SDS in PBS at 40 °C overnight. At each reagent replacement step, the gel beads were washed three times with DPBS and subsequently resuspended in the next solution. Following lysis, gel beads were washed with DPBS five times and the supernatant was removed. Then, the beads were suspended in Buffer D2 and subjected to multiple displacement amplification (MDA) using a REPLI-g Single Cell Kit (QIAGEN).

Following WGA at 30 °C for 2 h, gel beads were washed three times with 500 μL DPBS. Thereafter, beads were stained with 1× SYBR Green (Thermo Fisher Scientific) in DPBS. Following confirmation of DNA amplification by the presence of green fluorescence in the gel, fluorescence-positive beads were sorted into 0.8 μL DPBS in 96-well plates using a FACSMelody cell sorter (BD Bioscience) equipped with a 488-nm excitation laser. Following droplet sorting, 96-well plates proceeded to the second round of WGA or were stored at −30 °C.

Following gel bead collection in 96-well plates, second-round MDA was performed with the REPLI-g Single Cell Kit. Buffer D2 (0.6 μL) was added to each well and incubated at 65 °C for 10 min. Thereafter, 8.6 μL of MDA mixture was added and incubated at 30 °C for 120 min. The MDA reaction was terminated by heating at 65 °C for 3 min. Following second-round amplification, master library plates of SAGs were prepared. For quality control, aliquots of SAGs were transferred to replica plates for DNA yield quantification using a Qubit dsDNA High Sensitivity Assay Kit (Thermo Fisher Scientific). For sequencing analysis, sequencing SAG libraries were prepared from the second-round MDA products using QIAseq FX DNA Library Kit (QIAGEN). Ligation adaptors were modified to TruSeq™– Compatible Full-length Adapters UDI (Integrated DNA Technologies). Each SAG library was sequenced using an Illumina HiSeq 2 × 150 bp configuration (Macrogen).

### 16S rDNA sequencing

To confirm amplification from single-cell genomes and to identify the taxonomy from the mock community sample, 16S rRNA gene fragments V3–V4 were amplified with 341F and 806R primers (Forward, 5′-TCGTCGGCAGCGTCAGATGTGTATAAGAGACAGCCTACGGGNGGCWGCAG-3′; reverse, 5′-GTCTCGTGGGCTCGGAGATGTGTATAAGAGACAGGACTACHVGGGTATCTAATCC-3′) and sequenced by Sanger sequencing from SAGs obtained by SAG-gel. Following taxonomy identification with BLAST, every two to four species for mock communities were selected for whole-genome sequencing.

### Metagenome sequencing

Total DNA was extracted from mock samples according to the International Human Microbiota Standard protocol Q[36]. The DNeasy Power Soil Pro Kit (QIAGEN) was used for total DNA extraction from fecal and skin swab samples. Metagenomic sequencing libraries were constructed from extracted DNA samples with 10-μL (1/5 volume) reactions of the QIAseq FX DNA Library Kit. Each metagenomic sequencing library was sequenced using an Illumina HiSeq 2 × 150 bp configuration (Macrogen).

### Pre-processing and assembly of single-cell genomic and metagenomic sequence reads

SRs and MRs were individually processed for eliminating LQ reads by using fastp 0.20.1[37] with default options or bbduk.sh 38.79[38] (options: qtrim=r, trimq=10, minlength=40, maxns=1, minavgquality=15). Human genome contaminations were removed from SRs and MRs by mapping with bbmap.sh 38.79. SRs were assembled *de novo* using SPAdes 3.14.0 (options for SAG: --sc --careful --disable-rr --disable-gzip-output -t 4 -m 32), and contigs <1000 bp were excluded from subsequent analyses[39]. The MRs were assembled into contigs *de novo* using SPAdes 3.14.0 (options: --meta, -t 12, -m 96).

### Grouping of the same strain SAGs into CoSAG

SAGs with the completeness >10% in the mock community, 20% in the human microbiota sample, and contamination of <10% were selected with CheckM[40]. ANI was calculated for selected SAGs using fastANI 1.3[41]. The homology of common single-copy marker genes obtained using CheckM v1.1.2 taxonomy workflow (option: -nt --tab_table -t 16 domain Bacteria) was calculated by blastn 2.9.0+ with the default option. SAGs with ANI >95%, single-copy marker gene homology >99%, and tetra-nucleotide frequencies correlation >90% were identified in the same strain group. SRs from one SAG were mapped to other SAGs in the same group using MINIMAP2 2.17 (options: -ax sr)[42]. According to the ccSAG procedure[18], potential chimeras that partially aligned were split into aligned and unaligned fragments. The short fragments (<20 bp) were discarded. Clean and chimera-removed reads were obtained using cycles of cross-reference mapping and chimera splitting for each sample in the same group. Quality controlled reads from the same group were co-assembled *de novo* as CoSAG using SPAdes (options: --sc --careful --disable-rr --disable-gzip-output -t 4 -m 32).

### SAG-guided binning of metagenome contigs

The MAs were individually mapped against the strain-specific nrSAG contig using BWA 0.7.17 with the default option[43]. MA contigs that showed >99% identity (>200bp) to nrSAG contigs were extracted to construct sgBins by assignment based on nrSAG taxa.

### Merging of nrSAG and sgBin

For the two sets of assemblies, nrSAG and sgBin, CheckM was performed to measure their completeness, and the assembly with higher completeness was defined as the master and that with lower completeness as the slave. Slave assemblies were processed using SeqKit[44], and contigs <10000 bp were removed. Master and slave assemblies were merged using HaploMerger2_20180603[45] to create sgMAGs or mgSAGs. Thereafter, MAGs were reconstructed by using the DAS-tool from MAs that were unclassified as sgBin.

### Conventional MAG binning

For comparison of MAG quality, multiple binning of metagenomic contigs were conducted with conventional binners including CONCOCT 1.0.0[10], MaxBin 2 2.2.6[12], and MetaBAT 2 2.12.1[11] with default options. To refine the binning results obtained using these three different methods, DAS_Tool 1.1.2[13] was used with default options.

### Gene prediction, taxonomy identification, and plasmid detection

CDS, rRNAs, and tRNAs were extracted from all SAGs or MAGs by Prokka 1.14.6[46] (option: --rawproduct --mincontiglen 200). 16S and 23S rRNA genes with lengths ≥700 and 1100 bp, respectively, were detected. Taxonomy identification was performed using GTDB-Tk 1.3.0[47] with the default option, using the Release95 database. PlasClass[48] was used for detecting plasmids.

### Quality assessment of draft genomes from the mock community

In the mock community sample analysis, ANIs of each draft genome (sgMAG and mgSAG) for the closest reference genome were calculated with fastANI 1.3. The closest taxa with ≥ 99.5% ANI was assigned to each draft genome.

The quality of all obtained SAGs and MAGs were evaluated using QUAST v.5.0.2 (default option)[49], ChcekM v1.1.2 lineage workflow (option: --nt --tab_table -t 16), and identification of 5S, 16S, and 23S rRNA. To assess the accuracy of procured draft genomes in mock community samples, draft genomes were individually mapped to the corresponding taxa reference genome using MINIMAP2 2.17 with default options. The mapping results were converted to the pileup textual format using samtools 1.9[50], and the genomic coverage L for the reference genome was calculated using the following equation.

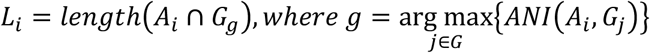

where A_i_represents the i^th^ draft genome. G and G_j_represent the set of reference genomes and the j^th^ reference genome of the set, respectively. G_g_ represents the corresponding reference genome against A_i_. When the reference genome is G_g_and the draft genome is A_i_, precision (P), recall(R), and F value (F_1_score) of the reference genome were calculated using the following equations.

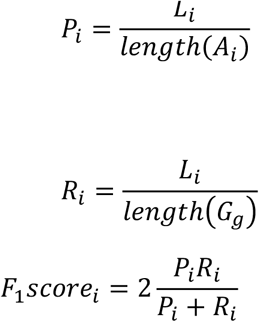

## Declarations

### Ethics approval and consent to participate

Studies with human subjects were approved by the School of Science and Engineering at Waseda University (No. 2018-323 and No. 2019-381). The subjects gave their written informed consent prior to sample collection.

## Consent for publication

Not applicable.

## Availability of data and material

MetaSAG is available from https://github.com/kojiari/metasag. Sequencing data has been deposited in the NCBI database under the BioProject PRJNA692334 (see Additional file 6: Table S5 for details).

## Competing interests

MH and HT are shareholders in bitBiome, Inc., which provides single-cell genomics service using SAG-gel workflow as bit-MAP. MH is a founder of bitBiome, Inc. KA, TS, TY, TE, and AM are employed at bitBiome, Inc. KA, KI, MK, HT, and MH are inventors on patent applications submitted by bitBiome, Inc. covering the technique for integration of metagenome and single-cell genome data.

## Funding

This work was supported by Tokyo Metropolitan Small and Medium Enterprise Support Center.

## Authors’ contributions

KA, HT, and MH conceived and designed the experiments. KA, KI, MK, and MH developed MetaSAG. TS, TY, TE, and AM conducted genomics experiments and collected the data. KA and KI conducted bioinformatic analysis of the metagenomic data and single-cell genomic data. KA and MH wrote the manuscript. All authors read and approved the final manuscript.

## Acknowledgements

The super-computing resource was provided by the Human Genome Center (University of Tokyo). Figure 1 was created with BioRender.com.

## Notes

### Summary of Updates

Title changed; Software name changed; Figures updated; Supplemental files updated.

