## Additional file 5 for "Recovery of high-quality assembled genomes via metagenome binning guided with single-cell amplified genomes"

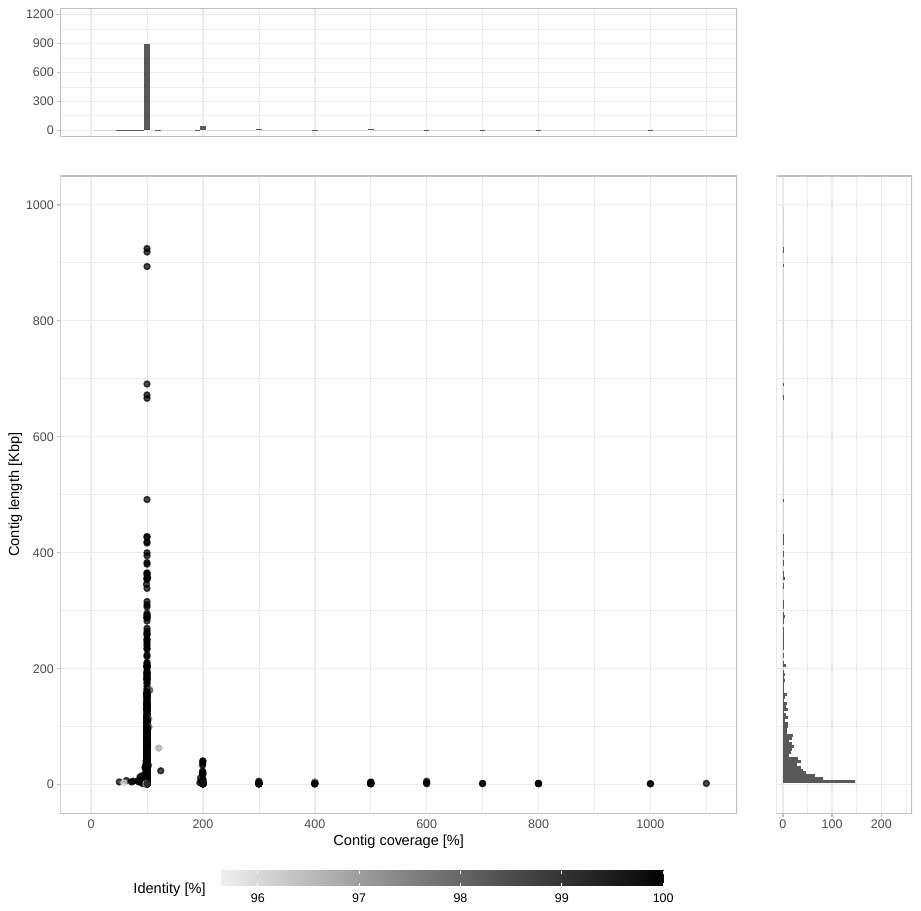

**Fig. S1 Distribution of coverage and length of contigs mapped on bacterial genomes contained in a cell mock community.** Regarding contig assignment, 91.1% of the total contigs were fully mapped to the single reference genome with ≥99% homology, while the contigs with the coverage of ≥200% were mapped to the repeating sequence part of the reference genomes. The majority of them were <5k bases in length.

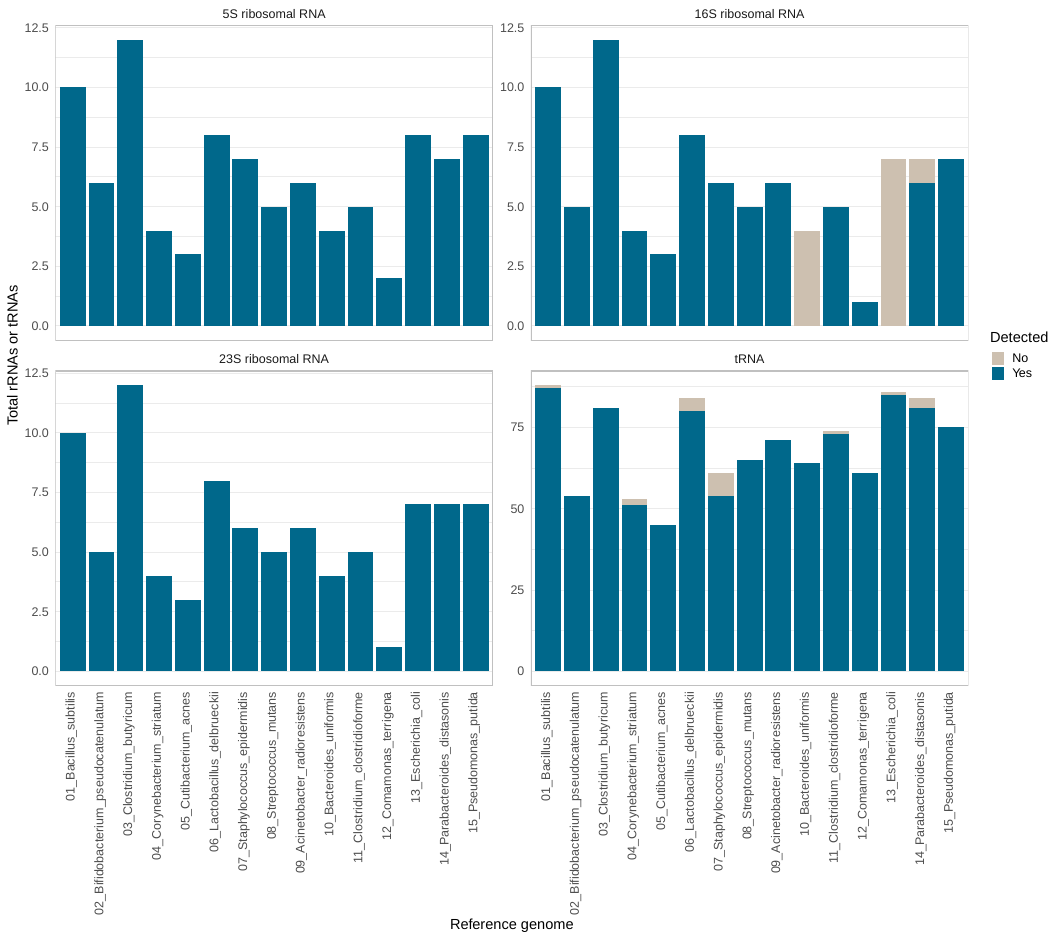

**Fig. S2 Number of rRNA genes and tRNA genes in metagenome assembled contigs (MAs) of 15 bacteria of a microbial community before binning.**

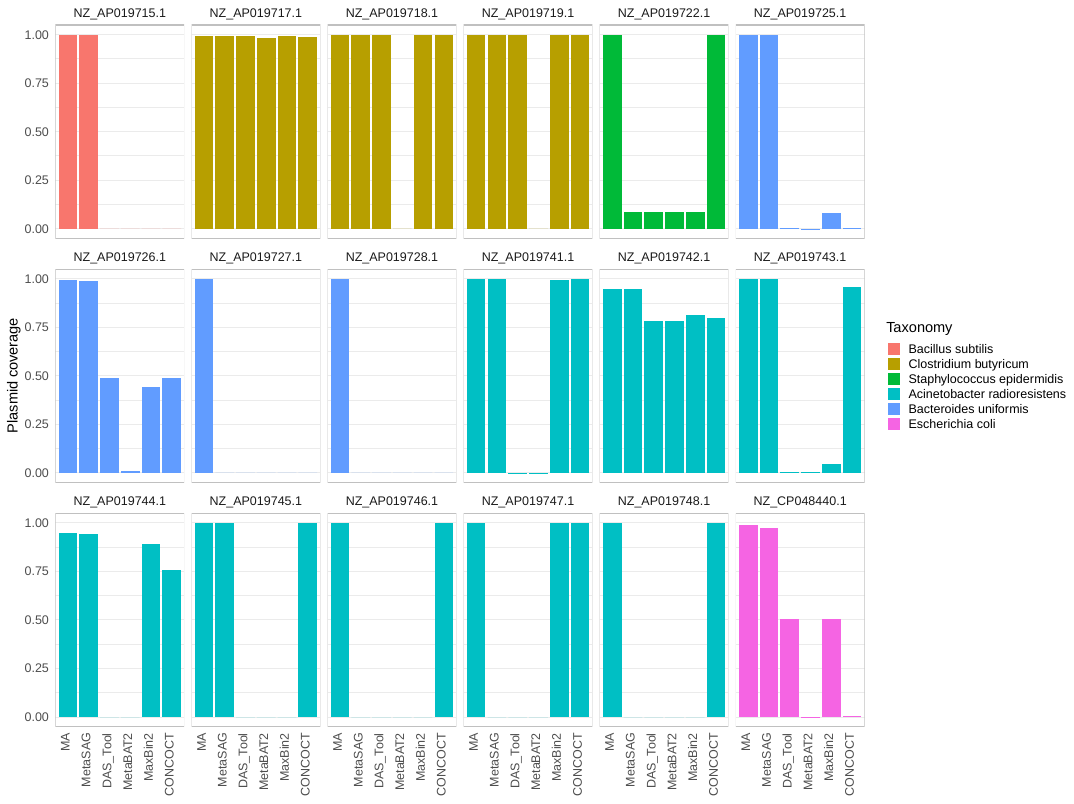

**Fig. S3 The performance of metagenomic assembly and binning in recovery of plasmid sequences.** Each bar shows the coverage of plasmid estimated from sequences in metagenome assembled contigs (MAs) and bins obtained with single-cell genome-guided binning of metagenomic assemblies (MetaSAG) and the other four binners. All data were collected from six plasmid-containing bacteria in a cell mock community.

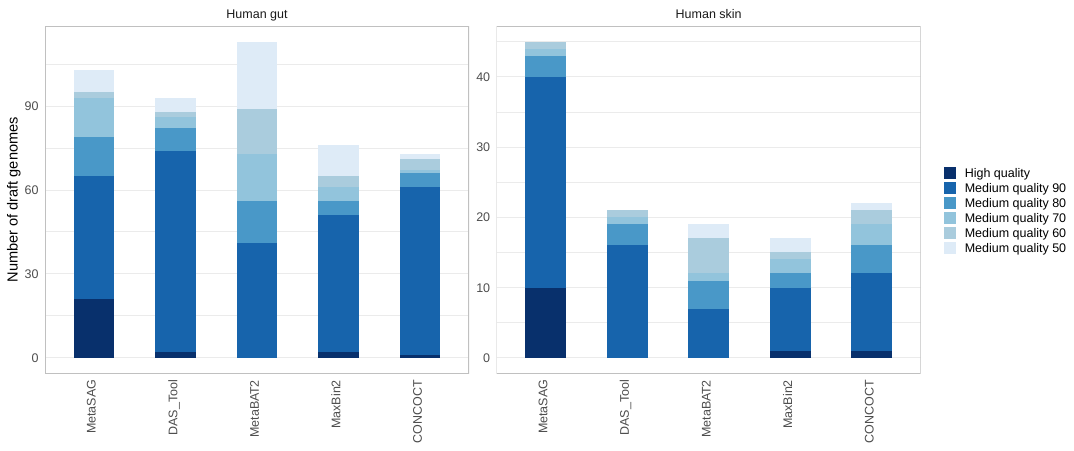

**Fig. S4 Draft genomes reconstructed from human microbiota samples by single-cell genome-guided binning of metagenomic assemblies (MetaSAG) and other binners with doubled metagenomic data.** All data were collected from three human fecal samples and three human skin swab samples. Single-cell genomic sequence reads (8.7 Gb) and metagenomic sequence reads (6.0 Gb) were used in MetaSAG. Doubled metagenomic sequence reads (approximately 14.9 Gb) were used in the other four binners.

**Tables**

Table S1 Cell mock community reference genome

Table S2 Sequence reads obtained from single-cell amplified genomes (SAGs) and metagenome

Table S3 Single-cell amplified genome (SAG) to composite single-cell amplified genome (CoSAG)

Table S4 Assembly quality of composite single-cell amplified genomes (CoSAGs) of cell mock community

Table S5 High-quality (HQ) and medium-quality (MQ) draft genomes constructed using single-cell genome-guided binning of metagenomic assemblies (MetaSAG)
